# Rapid cortical mapping with cross-participant encoding models

**DOI:** 10.64898/2026.03.26.714320

**Authors:** Jerry Tang, Alexander G. Huth

## Abstract

Voxelwise encoding models trained on functional MRI data can produce detailed maps of cortical organization. However, voxelwise encoding models must be trained on many hours of brain responses from each participant, limiting clinical applications. In this study, we introduce a cross-participant modeling framework for rapid cortical mapping. In this framework, voxelwise encoding models are trained on many hours of brain responses from previously scanned reference participants, and then transferred to a new participant by aligning brain responses using a small set of stimuli. We evaluated cross-participant encoding models on linguistic semantic mapping, non-linguistic semantic mapping, and auditory mapping. In each case we found that cross-participant encoding models had more accurate selectivity estimates and prediction performance than within-participant encoding models trained on the same amount of data from the new participant. We also found that cross-participant encoding models improved with the amount of data from each reference participant and the number of reference participants. These results demonstrate that cross-participant modeling can substantially reduce the amount of data required for accurate cortical mapping, which may facilitate new clinical applications of functional neuroimaging.

## 1. Introduction

A central goal of neuroscience is to map the information that is represented in each brain region. Recent studies have produced detailed cortical maps by training voxelwise encoding models on naturalistic neuroimaging data (Naselaris et al., 2011). In this approach, functional MRI (fMRI) is used to measure brain activity while participants perform naturalistic tasks such as viewing pictures (Allen et al., 2022; Eickenberg et al., 2017; Güçlü & van Gerven, 2015; Hebart et al., 2023; Kay, Naselaris, et al., 2008), watching movies (J. Chen et al., 2017; Hanke et al., 2014; Huth et al., 2012; Khosla et al., 2021; Nishimoto et al., 2011; Seeliger et al., 2019; Visconti di Oleggio Castello et al., 2020), and reading or listening to stories (de Heer et al., 2017; Deniz et al., 2019; Huth et al., 2016; Jain et al., 2020; Millet et al., 2022; Nastase et al., 2021; Schrimpf et al., 2021; Vaidya et al., 2022; Wehbe et al., 2014). These ecologically valid experiments generate datasets that capture complex relationships between the stimuli and brain responses (Deniz et al., 2023; Hamilton and Huth, 2018; Nastase et al., 2020). To model these data, quantitative features are extracted from the stimuli to capture different hypotheses for how stimulus information is represented in the brain. For instance, different stages of language processing can be modeled using spectrograms, phonemes, part-of-speech tags, and distributed semantic features (de Heer et al., 2017; LeBel et al., 2021; Wehbe et al., 2014). Finally, linear regression is used to learn a set of encoding model weights that predict how each voxel responds to the stimulus features.

Previous studies have analyzed encoding model weights to determine the stimuli that each region is selective for (Huth et al., 2016; Jain et al., 2023) and define clusters of regions with similar selectivity (LeBel et al., 2021; Meschke et al., 2023). Other studies have used encoding model performance to characterize the features that are encoded in each region (de Heer et al., 2017; Eickenberg et al., 2017; Güçlü & van Gerven, 2015; LeBel et al., 2021; Wehbe et al., 2014) and identify regions that perform similar functions across different tasks (C. Chen et al., 2024; Deniz et al., 2019; Popham et al., 2021; Tang, Du, et al., 2023). All of these analyses produce detailed cortical maps in each participant. These cortical maps could potentially be used to monitor cognitive status (Fischer et al., 2025; Sperling, 2011; Voineskos et al., 2024), plan surgical interventions (Silva et al., 2018), localize surgical implant and neurostimulation sites (Leinders et al., 2023; Richardson et al., 2015; Verwoert et al., 2025), and develop neurofeedback targets (Taschereau-Dumouchel et al., 2018). However, a major barrier for clinical applications is that training an encoding model for an individual participant currently requires many hours of brain responses from that participant (LeBel et al., 2023).

When only a limited amount of data can be collected from a new *goal participant*, one potential solution is to incorporate data from other *reference participants*. Previous studies have shown that the set of specialized functional regions is highly consistent across individuals, even if the precise anatomical arrangement of the regions is not (Braga & Buckner, 2017; P.-H. Chen et al., 2015; Fedorenko et al., 2010; Haxby et al., 2011; Huth et al., 2016; Kanwisher et al., 1997). Functional correspondences between brain regions in different participants can be identified using responses to shared stimuli. If a region in a reference participant has similar responses to a region in a goal participant, it indicates that the two regions are selective for similar information. Models for the region in the reference participant can then be transferred to the region in the goal participant. Functional alignment has previously been used to transfer decoding models across participants (Ho et al., 2023; Tang & Huth, 2025; Yamada et al., 2015) and identify representational spaces that are shared across participants (Haxby et al., 2011). However, it is unclear whether functional alignment can be also used to transfer detailed cortical maps.

Here we introduce a framework for transferring encoding models across participants in order to perform cortical mapping with limited data. In this framework, we record brain responses from reference participants while they listen to many hours of narrative stories (**Figure 1A**). We use these data to train a reference encoding model that takes quantitative stimulus features and predicts how the reference participants would respond. To transfer the model to a new goal participant, we record brain responses from both the reference participants and the goal participant using a small, common set of alignment stimuli (**Figure 1B**). We use these data to train a *cross-participant converter* that takes brain responses from the reference participants and predicts how the goal participant would respond to the same stimulus. Finally, we create a cross-participant encoding model by projecting the reference encoding model through the cross-participant converter (**Figure 1C**). This cross-participant encoding model takes quantitative stimulus features and predicts how the goal participant would respond. Our results demonstrate that cross-participant modeling is an accurate alternative to within-participant modeling that requires far less data from the goal participant. We have developed an open-source Python package to support future studies and applications.

**Figure 1.**
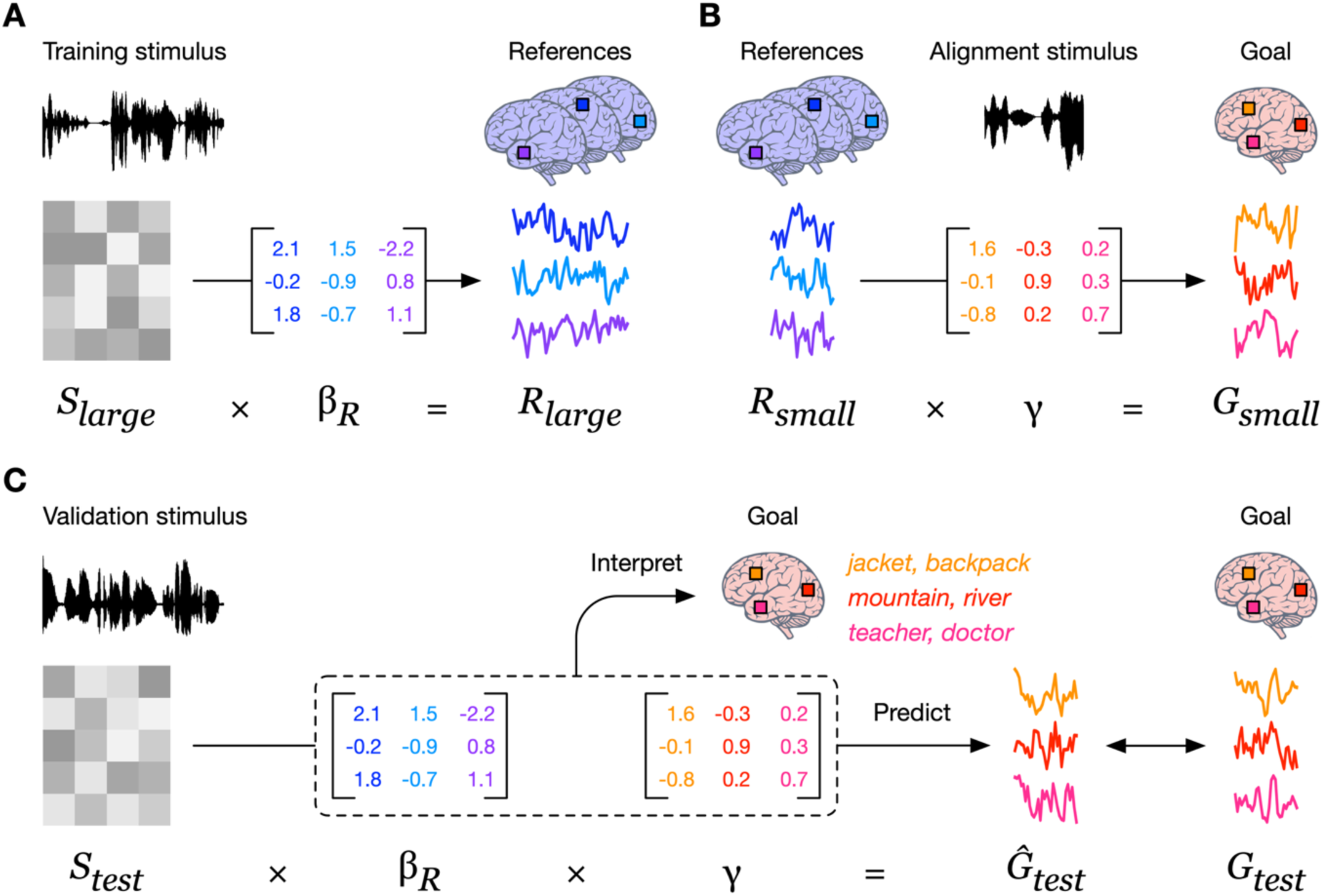
Transferring encoding models across participants. (**A**) Brain responses to a large set of narrative stories are recorded from reference participants. A reference encoding model *β*_*R*_ is trained to predict the reference participant responses based on quantitative stimulus features. (**B**) Brain responses to a shared set of narrative stories or silent movies are recorded from the reference participants and the goal participant. A linear cross-participant converter *γ* is trained to predict the goal participant responses based on the reference participant responses. (**C**) Multiplying the reference encoding model *β*_*R*_ by the cross-participant converter *γ* produces a cross-participant encoding model that predicts goal participant responses based on quantitative stimulus features. The cross-participant encoding model weights *β*_*R*_*γ* can be interpreted to characterize the selectivity of each brain region in the goal participant. The cross-participant encoding model can also be evaluated by predicting goal participant responses to held-out stories and taking the linear correlation between the predicted and actual response time-courses.

## 2. Methods

### 2.1 Cross-participant modeling framework

For many clinical applications, only a limited amount of data can be collected from a new *goal participant*. The goal participant is presented with a small stimulus set which evokes brain responses *G*_*small*_ ∈ *ℝ*^*n*×*g*^ where *n* is the number of timepoints and *g* is the number of voxels in the goal participant’s brain. The small stimulus set can be represented using the quantitative features *X*_*small*_ ∈ *ℝ*^*n*×*f*^ where *f* is the number of features. Using the typical within-participant voxelwise modeling approach, the encoding model *β*_*G*_ ∈ *ℝ*^*f*×*g*^ predicts brain responses from the stimulus features. The encoding model is defined as *G*_*small*_ = *X*_*small*_*β*_*G*_ and is estimated using regularized linear regression as *β*_*G*_ = *X*_*small*_^+^*G*_*small*_, where *X*^+^ is a pseudoinverse of *X*. We typically use ridge regression, which has a single free parameter that controls the degree of regularization (Hoerl & Kennard, 1970). This regularization parameter is optimized by repeating a cross-validation procedure. In each iteration of this procedure, approximately a fifth of the timepoints are removed from the model training dataset and reserved for validation. Then, the model weights are estimated on the remaining timepoints for each of 10 possible regularization coefficients (log spaced between 10 and 1,000). These weights are used to predict responses for the reserved timepoints, and then *R^2^* is computed between actual and predicted responses. The regularization coefficient is chosen as the value that leads to the best performance, averaged across voxels and iterations, on the reserved timepoints.

In cross-participant modeling, a larger amount of data can also be collected from a set of reference participants. The reference participants are presented with a large stimulus set which evokes brain responses *R*_*large*_ ∈ *ℝ*^*m*×*r*^ where *m* ≫ *n* is the number of timepoints and *r* is the number of voxels across all reference participants. Since only a fraction of cortical voxels will be selective for the stimulus features, *r* is restricted to the 10,000 best performing voxels from each reference participant. The large stimulus set is represented using the quantitative features *X*_*large*_ ∈ *ℝ*^*m*×*f*^. The reference encoding model *β*_*R*_ ∈ *ℝ*^*f*×*r*^ is defined as *R*_*large*_ = *X*_*large*_*β*_*R*_ and is again estimated with regularized linear regression as *β*_*R*_ = *X*_*large*_^+^*R*_*large*_.

The reference encoding model is transferred to the goal participant by training a cross-participant converter. To perform functional alignment, the reference participants are presented with the same small stimulus set that was used for the goal participant, evoking brain responses *R*_*small*_ ∈ *ℝ*^*n*×*r*^. The cross-participant converter *γ* ∈ *ℝ*^*r*×*g*^ is defined as *G*_*small*_ = *R*_*small*_*γ* and is estimated with regularized linear regression as *γ* = *R*_*small*_^+^*G*_*small*_. The cross-participant converter predicts how the activity in each goal participant voxel corresponds to the activity in a population of reference participant voxels. The reference encoding model *β*_*R*_ is a mapping from stimulus features to reference participant voxels, and the cross-participant converter *γ* is a mapping from reference participant voxels to goal participant voxels. These two mappings can thus be composed to give the cross-participant encoding model *β*^∗^_*small*_ = *β*_*R*_*γ*, which is a mapping from stimulus features to goal participant voxels.

To give an intuition for why cross-participant encoding models are effective, we can perform some substitutions. The cross-participant model can be written as *β*^∗^_*small*_ = *β*_*R*_*γ* = (*X*_*large*_^+^*R*_*large*_)(*R*_*small*_^+^*G*_*small*_) which resembles the within-participant model *X*_*small*_^+^*G*_*small*_ where *X*_*small*_^+^ is replaced by *X*_*large*_^+^*R*_*large*_*R*_*small*_^+^. This can be interpreted as expressing each time-point in the small stimulus set as a linear combination of the time-points of the large stimulus set. The interpolation operator *R*_*large*_*R*_*small*_^+^ enables this by comparing the reference participant responses to the small and large stimulus sets. As a result, cross-participant modeling can also be viewed as using data from the reference participants to learn stimulus representations that more closely reflect the representational space of reference participant brain responses.

Analyses were performed using custom software written in Python, making heavy use of NumPy (Harris et al., 2020) and pycortex (Gao et al., 2015).

### 2.2 Encoding model validation

For cross-participant encoding models to be useful, they should be similar to encoding models trained on large amounts of data from the goal participant. Previous studies have found that encoding model weights stabilize after several hours of training data (LeBel et al., 2023). Thus, we evaluated cross-participant encoding models by comparing them to *ceiling encoding models* trained on 5 hours of brain responses from the goal participant. The ceiling training dataset was kept separate from the reference training dataset and the small functional alignment dataset. We characterized the selectivity of each voxel by constructing a large set of *n* stimuli and using the encoding models to simulate how each voxel responds to every stimulus (Huth et al., 2016). One way to quantify whether two models have similar selectivity is using a *word overlap score* where we identify the top *k* stimuli for each model and compute the fraction that overlap. Chance level 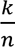 is given by the expected overlap between two random subsets of *k* elements from a set of *n* elements. Another way to quantify whether two models have similar selectivity is using a *selectivity correlation score* where we compute the rank correlation between the simulated responses to all *n* stimuli.

For cross-participant encoding models to be useful, they should also be able to generalize to new stimuli. We used encoding models to predict brain responses to a test story that was held out from training. We evaluated generalization for each voxel using a *prediction performance score* where we compute the linear correlation between the predicted and actual response time-courses.

### 2.3 MRI data collection

Data were collected from four participants: S1 (male, age 38 years at time of most recent scan), S2 (male, age 25 years), S3 (male, age 22 years), and S4 (female, age 27 years). All participants were healthy and had normal hearing, and normal or corrected-to-normal vision. Written informed consent was obtained from all participants. The experimental protocol was approved by the Institutional Review Board at the University of Texas at Austin.

MRI data were collected on a 3T Siemens Skyra scanner, a 3T Siemens Vida scanner, and a 3T Siemens Prisma scanner at the UT Austin Biomedical Imaging Center using a 64-channel Siemens volume coil. Functional data were collected using gradient echo EPI with repetition time (TR) = 2.00 s, echo time (TE) = 30.8 ms, flip angle = 71, multi-band factor (simultaneous multi-slice) = 2, voxel size = 2.6mm x 2.6mm x 2.6mm (slice thickness = 2.6mm), matrix size = (84, 84), and field of view = 220 mm. Anatomical data for all participants except S1 were collected using a T1-weighted multiecho MP-RAGE sequence on the same 3T Siemens Skyra scanner with voxel size = 1mm x 1mm x 1mm following the Freesurfer morphometry protocol. Anatomical data for participant S1 were collected on a 3T Siemens TIM Trio scanner at the UC Berkeley Brain Imaging Center with a 32-channel Siemens volume coil using the same sequence.

### 2.4 MRI experiments

10 hours of data were collected while each participant listened to autobiographical narrative stories from *The Moth Radio Hour* and *Modern Love* (LeBel et al., 2023; Tang, LeBel, et al., 2023). Each story was played during a separate fMRI scan with a buffer of 10 s of silence before and after the story. Stories were played over Sensimetrics S14 in-ear piezoelectric headphones. The audio for each stimulus was filtered to correct for frequency response and phase errors induced by the headphones using calibration data provided by Sensimetrics and custom Python code (https://github.com/alexhuth/sensimetrics_filter). Each story was manually transcribed by one listener. Certain sounds (for example, laughter and breathing) were marked to improve the accuracy of the automated alignment. The Penn Phonetics Lab Forced Aligner (P2FA) was used to automatically align the audio to the transcript (Yuan & Liberman, 2008), and Praat was used to check and correct each aligned transcript manually (Boersma & Weenink, 2014).

1 hour of data was collected while each participant watched movie clips from *Pixar Animation Studios* and *The Blender Foundation* (Tang & Huth, 2025). The movie clips were self-contained and almost entirely devoid of language. The original high-definition movie clips were cropped and downsampled to 727 x 409 pixels. Each movie clip was played without sound during a separate fMRI scan with a 10 s black screen buffer before and after the movie clip.

A repeated test dataset was collected while each participant listened to 5 repeats of the 10 m test story ‘‘Where There’s Smoke’’ by Jenifer Hixson from *The Moth Radio Hour*. Each repeat of the test story was played during a single fMRI scan with a buffer of 10 s of silence before and after the story. The 5 repeats were collected across multiple scanning sessions. The test story was held out from model training and used to compute prediction performance scores.

To evaluate the cross-participant modeling approach, we treated each participant in turn as the goal participant and the remaining participants as the reference participants. The reference encoding models were trained on the first 5 hours of brain responses to stories. The functional alignment stories were chosen from a subset of 20 stories from the first 5 hours that are each 10- 14 m. The functional alignment movies were chosen from a set of 8 movies that are each 4-6 m. Results were averaged across 5 permutations of the functional alignment stimuli to control for differences in data quality. The ceiling encoding models were trained on the last 5 hours of brain responses to stories. This avoids circularity by ensuring that the ceiling training data are separate from the reference training data and the functional alignment data.

### 2.5 MRI preprocessing

Each functional run was motion-corrected using the FMRIB Linear Image Registration Tool (FLIRT) from FSL 5.0 (Jenkinson & Smith, 2001). All volumes in the run were then averaged to obtain a high quality template volume. FLIRT was used to align the template volume for each run to the overall template, which was chosen to be the template for the first functional run for each participant. These automatic alignments were manually checked. Low-frequency voxel response drift was subtracted from the signal. The mean response for each voxel was then subtracted and the remaining response was scaled to have unit variance. For responses to stories, low-frequency voxel response drift was identified using a 2nd order Savitsky-Golay filter with a 120 second window. For responses to movies, low-frequency voxel response drift was identified using a Legendre polynomial of degree 3, since the movie scans were shorter than the story scans (Kay, David, et al., 2008).

Cortical surface meshes were generated from the T1-weighted anatomical scans using Freesurfer (Dale et al., 1999). Before surface reconstruction, anatomical surface segmentations were hand-checked and corrected. Blender was used to remove the corpus callosum and make relaxation cuts for flattening. Functional images were aligned to the cortical surface using boundary based registration (BBR) implemented in FSL. These alignments were manually checked for accuracy and adjustments were made as necessary. Flatmaps were created by projecting the values for each voxel onto the cortical surface using the ‘‘nearest’’ scheme in pycortex (Gao et al., 2015). This projection finds the location of each pixel in the flatmap in 3D space and assigns that pixel the associated value.

### 2.6 Linguistic feature spaces

Story comprehension requires mapping from sounds to meanings. Different types of quantitative features can be extracted from the story stimuli to study different stages of language processing.

We modeled high-level semantic processing using a distributional feature space derived from word co-occurrence statistics (Huth et al., 2016). We constructed a 10,470 word lexicon from the union of the set of all words appearing in the first 2 story sessions and the 10,000 most common words in a large text corpus. We then selected 985 basis words from Wikipedia’s *List of 1000 Basic Words* (contrary to the title, this list contained only 985 unique words at the time it was accessed). This basis set was selected because it consists of common words that span a very broad range of topics. The text corpus used to construct this feature space includes the transcripts of 13 stories from *The Moth Radio Hour* (including 10 used as stimuli in this experiment), 604 popular books, 2,405,569 Wikipedia pages, and 36,333,459 user comments scraped from reddit.com. In total, the 10,470 words in our lexicon appeared 1,548,774,960 times in this corpus. Next, we constructed a word co-occurrence matrix *M* with 985 rows and 10,470 columns. Iterating through the text corpus, we added 1 to *M*_*ij*_ each time word *j* appeared within 15 words of basis word *i*. A window size of 15 was selected to be large enough to suppress syntactic effects but no larger. Once the word co-occurrence matrix was complete, we log-transformed the counts, replacing *M*_*ij*_ with *log* (*1* + *M*_*ij*_). Next, each row of *M* was *z*-scored to correct for differences in basis word frequency, and then each column of *M* was *z*-scored to correct for word frequency. At the end of this process, each column of *M* contains a 985- dimensional semantic vector representing one word in the lexicon.

We modeled lower-level auditory processing using spectral and articulatory feature spaces (LeBel et al., 2021). The spectral feature space was a mel-band spectrogram with frequencies ranging from ∼0 Hz to 8 kHz with 256 windows. The articulatory feature space was an n-hot feature space where each phoneme is assigned a 1 for each articulation that is required to produce the sound and a 0 for every other articulation for a total of 22 features for each phoneme.

### 2.7 Statistical testing

For each goal participant, we identified the voxels that are selective for each feature space by performing a permutation test on the repeated test dataset. We used the ceiling encoding model to predict brain responses to the test story, and computed the prediction performance score for each voxel as the linear correlation between the predicted and actual response time-courses. We then constructed a null distribution for each voxel by randomly resampling (with replacement) 10-TR blocks from the voxel’s actual response time-course. Resampling contiguous blocks preserves the auto-correlation structure of the voxel’s responses. We then computed the null prediction performance score as the linear correlation between the predicted and permuted response time-courses. Repeating this process for 1,000 trials provided a null distribution of prediction performance scores for each voxel. Significantly predicted voxels were identified as voxels with an observed prediction performance score that is significantly higher than its null distribution (*q*(FDR) < 0.05), correcting for multiple comparisons using the false discovery rate (Benjamini & Hochberg, 1995).

For each model, we averaged the word overlap, selectivity correlation, and prediction performance scores across significantly predicted voxels. This results in a cross-participant score and within-participant score for each goal participant. We compared the cross-participant scores to the within-participant scores using a two-sided paired t-test and corrected for multiple comparisons using the false discovery rate.

## 3 Results

### 3.1 Rapid semantic mapping

Semantic representations encode the meanings of concepts to provide an internal model of the world (Binder & Desai, 2011). Since semantic representations reflect the ideas that individuals are thinking about, semantic maps could be used to monitor cognitive status in neurological disorders such as dementia, schizophrenia, and disorders of consciousness (Fischer et al., 2025; Sperling, 2011; Voineskos et al., 2024). And since semantic representations are more widely distributed and less anatomically consistent than lower-level sensory and motor representations, semantic maps could also be important for localizing implant and stimulation sites for brain-computer interfaces that target higher-level cognitive processes (Jamali et al., 2024; Verwoert et al., 2025). Previous studies have modeled semantic representations in natural language using features derived from word co-occurrence statistics, which reflects the hypothesis that words which occur in similar contexts have similar meanings (Huth et al., 2016). While these models produce detailed cortical maps of semantic selectivity, they require many hours of training data due to the high dimensionality of semantic representations (LeBel et al., 2023).

Here we used small amounts of alignment data to transfer semantic encoding models from reference participants to the goal participant. We trained a reference encoding model on brain responses to 5 hours of stories. We transferred the reference encoding model to the goal participant using cross-participant converters trained on brain responses to subsets of the stories. To measure the benefits of cross-participant modeling, we trained within-participant models for the goal participant on brain responses to the same subsets of stories. We separately trained a ceiling encoding model for the goal participant on brain responses to 5 hours of different stories.

Previous studies have visualized semantic encoding models by projecting the weights onto the first 3 principal components of a group space (Deniz et al., 2019; Huth et al., 2016; Popham et al., 2021). Here we visualized the ceiling model trained on 5 hours of story data from the goal participant, a within-participant model trained on 24 minutes of story data from the goal participant, and a cross-participant model trained on 24 minutes of story data from the goal participant (**Figure 2A**). The ceiling model illustrates how semantic representations are actually organized across cortex for the goal participant. In posterior regions, concrete concepts are represented near visual cortex and somatosensory cortex, while abstract concepts are represented in medial parietal cortex, lateral temporal cortex, and the temporal-parietal junction. In anterior regions, concepts are tiled in intricate patches. The within-participant model provides a clear separation between concrete and abstract concepts but lacks the granularity of the ceiling model. For instance, regions that are mapped to *social*, *mental*, *violence*, and *person* concepts in the ceiling model are all mapped to *social* concepts in the within-participant model. By contrast, the cross-participant model is qualitatively far more precise and captures most of the details of the ceiling model. The main difference was in visual cortex, which is poorly predicted by language encoding models and might not encode shared representations during language processing (Huth et al., 2016; Tang & Huth, 2025).

**Figure 2.**
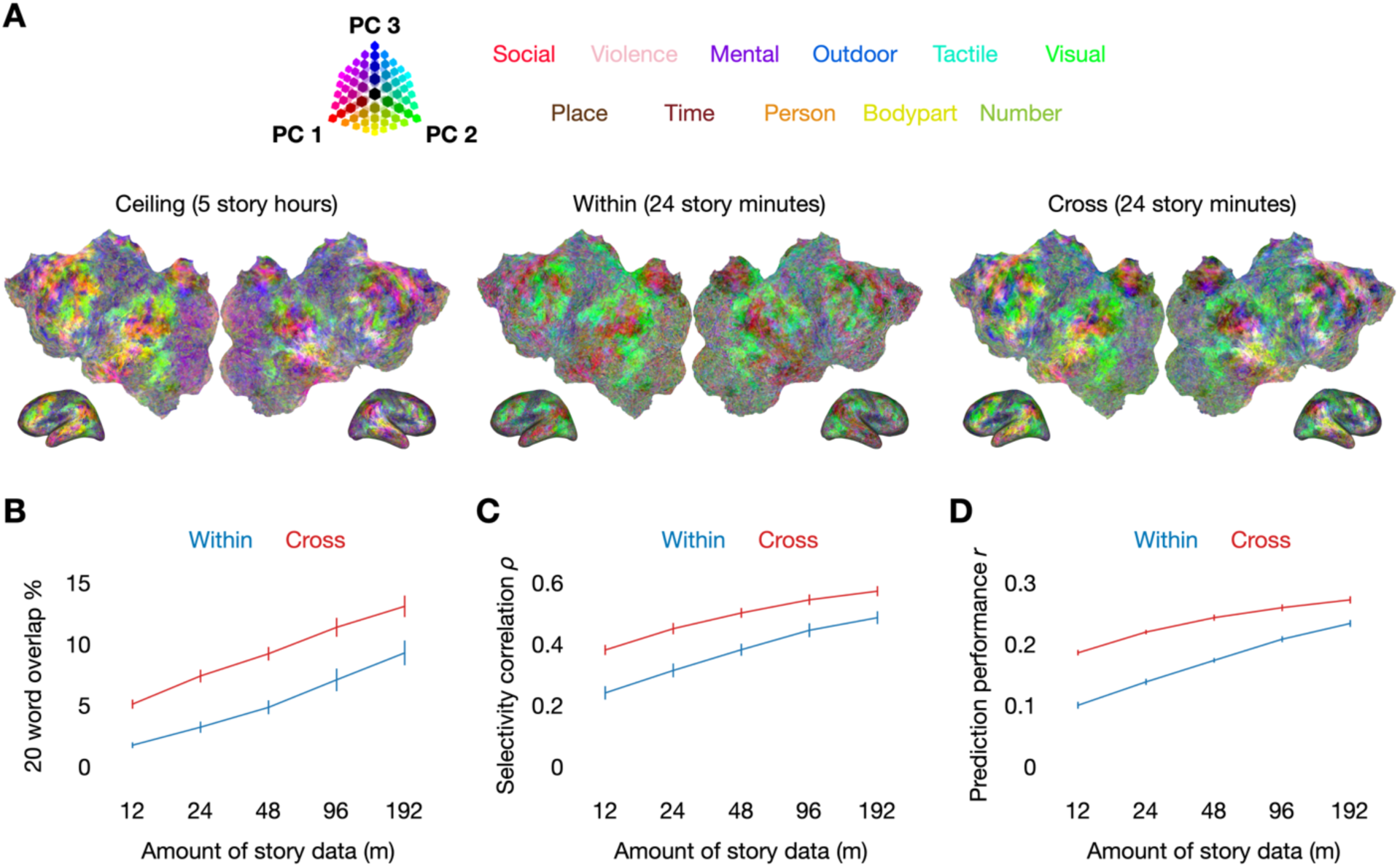
Rapid semantic mapping. Cross-participant and within-participant encoding models were trained on brain responses to narrative stories. The stimulus stories were represented using semantic features derived from word co-occurrence statistics. (**A**) Semantic selectivity was visualized by projecting encoding model weights onto the first 3 principal components of a group space. A cross-participant model trained on 24 minutes of story responses from a goal participant captured the same patterns of semantic selectivity as a ceiling encoding model trained on 5 hours of story responses from the goal participant. (**B**) Semantic selectivity was operationalized by simulating responses to 10,470 words and identifying the 20 words with the highest simulated responses. Cross-participant models had higher word overlap with the ceiling model than within- participant models trained on the same amount of data from the goal participant. (**C**) Semantic selectivity was alternatively operationalized by simulating responses to 10,470 words and ranking the words based on the simulated responses. Cross-participant models had higher selectivity correlation with the ceiling model than within-participant models trained on the same amount of data from the goal participant. (**D**) Prediction performance was quantified using the linear correlation between predicted and actual response time-courses for a held-out test story. Cross-participant models had higher prediction performance than within-participant models trained on the same amount of data from the goal participant.

To quantitatively assess whether cross-participant models can recover the same selectivity estimates as the ceiling model, we used the models to simulate brain responses to 10,470 words (Huth et al., 2016). For each significantly predicted voxel under the ceiling model, we computed the word overlap score between the top 20 words for the cross-participant models and the top 20 words for the ceiling model (**Figure 2B**). We found that cross-participant models could achieve over 10% word overlap with the ceiling model (chance level 0.19%), which demonstrates that they can be used to predict the concepts that are represented in each brain region. We also found that cross-participant models had a higher word overlap with the ceiling model than within-participant models trained on the same amount of data from the goal participant (*q*(FDR) < 0.05; two-sided paired t-test). While word overlap is an interpretable and functionally useful metric, it requires imposing an arbitrary cutoff on the number of words that are used to characterize selectivity. A more holistic alternative is to rank the simulated responses to all 10,470 words. We can then quantify the similarity between selectivity estimates by computing the selectivity correlation score between the ranks (**Figure 2C**). We found that cross-participant models had a higher selectivity correlation with the ceiling model than within-participant models trained on the same amount of data from the goal participant (*q*(FDR) < 0.05; two-sided paired t-test).

Finally we assessed how well cross-participant models can predict brain responses to a test story that was not used in model training or functional alignment. We quantified prediction performance for each voxel using the linear correlation between the predicted and actual response time-courses (**Figure 2D**). Across voxels that were significantly predicted by the ceiling model, we found that cross-participant models outperformed within-participant models trained on the same amount of data from the goal participant (*q*(FDR) < 0.05; two-sided paired t-test). While the advantage of cross-participant modeling decreased as we increased the amount of data from the goal participant, cross-participant models were still more effective when trained on over 3 hours of data from the goal participant.

### 3.2. Scaling with reference data

Our initial results show that using data from other reference participants can lead to more effective and accurate cortical maps when the amount of training data from a goal participant is limited. It may be possible to further improve the cross-participant maps by collecting more data from reference participants. Previous studies have found that within-participant encoding models scale with the amount of training data, so cross-participant encoding models may scale with the amount of training data from each reference participant (LeBel et al., 2023). Other studies have found that cross-participant decoding models scale with the number of reference participants, so cross-participant encoding models may also scale with the number of reference participants (Défossez et al., 2023; Ho et al., 2023; Tang & Huth, 2025). Here we assessed how the amount of data from each reference participant and the number of reference participants affect cross- participant modeling.

We first evaluated how the amount of data used to estimate encoding models in each reference participant affects cross-participant modeling. We trained reference participant models on brain responses from all of the reference participants, but varied the amount of data from each participant. We created the reference encoding model training datasets by including all of the functional alignment stories, and then adding 60 to 240 minutes of additional stories. We found that increasing the amount of data from each reference participant improved word overlap (**Figure 3A**), selectivity correlation (**Figure 3B**), and prediction performance (**Figure 3C**) for all amounts of data from the goal participant.

**Figure 3.**
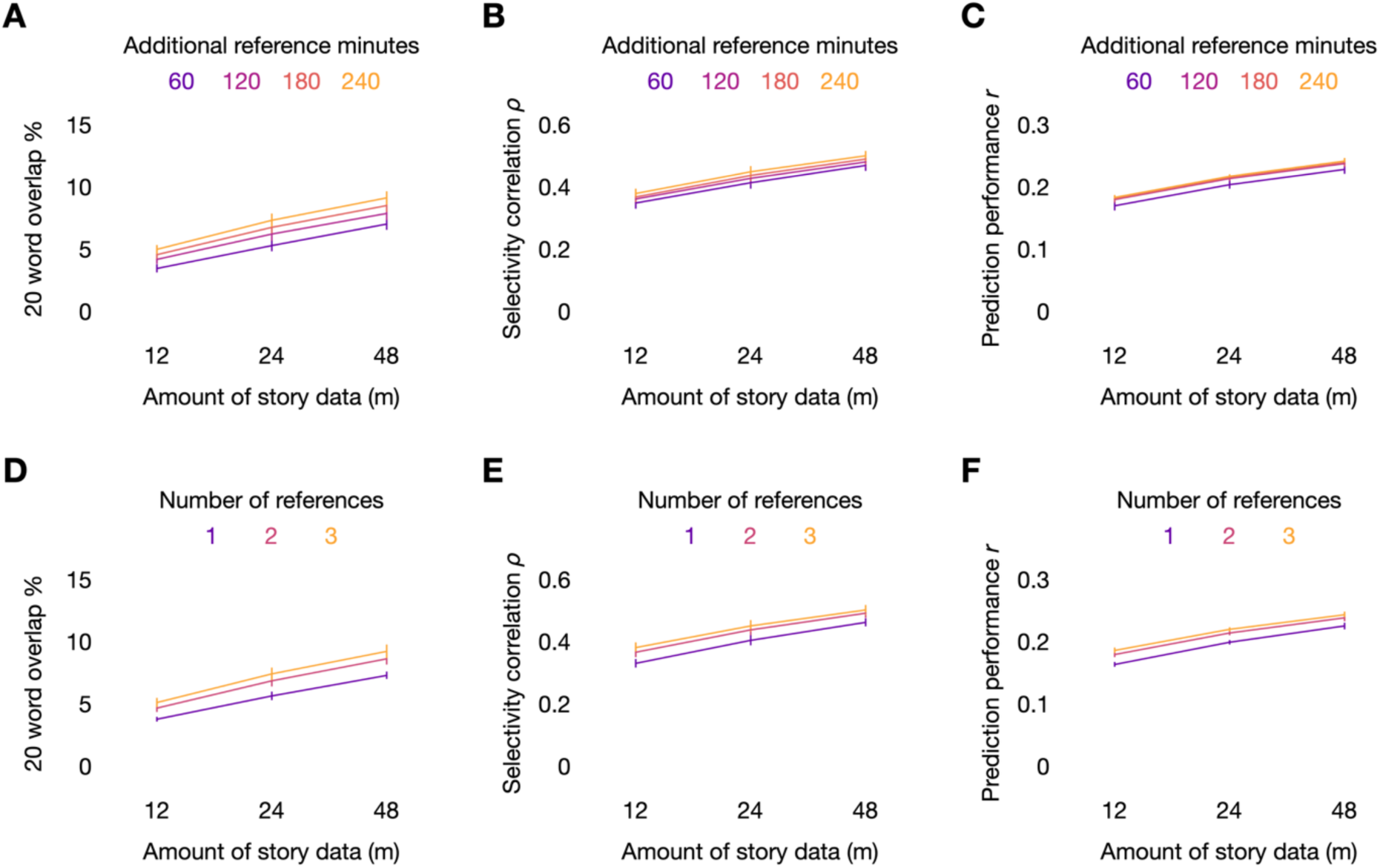
Scaling with reference data. Cross-participant encoding models were trained using different amounts of training data from each reference participant and different numbers of reference participants. The x-axes indicate the size of the functional alignment dataset. (**A**) Word overlap improved with the amount of data from each reference participant. (**B**) Selectivity correlation improved with the amount of data from each reference participant. (**C**) Prediction performance improved with the amount of data from each reference participant, but most of the improvement came between 60 and 120 minutes. (**D**) Word overlap improved with the number of reference participants, but more of the improvement came between 1 and 2 reference participants. (**E**) Selectivity correlation improved with the number of reference participants, but more of the improvement came between 1 and 2 reference participants. (**F**) Prediction performance improved with the number of reference participants, but more of the improvement came between 1 and 2 reference participants.

We next evaluated how the number of reference participants affects cross-participant modeling. We trained reference encoding models on brain responses to 5 hours of stories, but varied the number of reference participants. We found that increasing the number of reference participants also improved word overlap (**Figure 3D**), selectivity correlation (**Figure 3E**), and prediction performance (**Figure 3F**) for all amounts of data from the goal participant. These results demonstrate that cross-participant encoding models can be improved without increasing the amount of data from the goal participant.

### 3.3. Rapid cross-modal semantic mapping

Semantic representations are essential for language, but they are not exclusive to language. In particular, many brain regions are selective for the same concepts in language and vision (Devereux et al., 2013; Fairhall & Caramazza, 2013; Popham et al., 2021). Previous studies have trained semantic decoders on brain responses to narrative stories and transferred them across participants by aligning brain responses to silent movies (Tang & Huth, 2025). If semantic encoding models can also be transferred across participants using silent movies, it would provide a way to map semantic representations in children and individuals with language comprehension impairments.

Here we trained a reference encoding model on brain responses to 5 hours of stories using the same semantic features derived from word co-occurrence statistics. We transferred the reference encoding model to the goal participant using cross-participant converters trained on brain responses to silent movies. To measure the benefits of cross-participant modeling, we trained within-participant models for the goal participant using brain responses to the same movies. For these within-participant models, we transcribed official audio descriptions of the movies and extracted semantic features of the transcripts. We separately trained a ceiling encoding model for the goal participant on brain responses to 5 hours of different stories.

We visualized the ceiling model trained on 5 hours of story data from the goal participant, a within-participant model trained on 40 minutes of movie data from the goal participant, and a cross-participant model trained on 40 minutes of movie data from the goal participant (**Figure 4A**). While the within-participant encoding model can occasionally differentiate between concrete and abstract concepts, it does not reliably capture any of the details of the ceiling model. By contrast, the cross-participant model captures many details of the ceiling model. The main differences between the cross-participant model and the ceiling model are in visual cortex and auditory cortex, which reflects the fact that the models were trained on stimuli of different modalities (**Figure A1**). However, the cross-participant model and the ceiling model are similar in most other regions. This demonstrates that semantic concepts can be rapidly mapped purely based on brain responses to vision.

**Figure 4.**
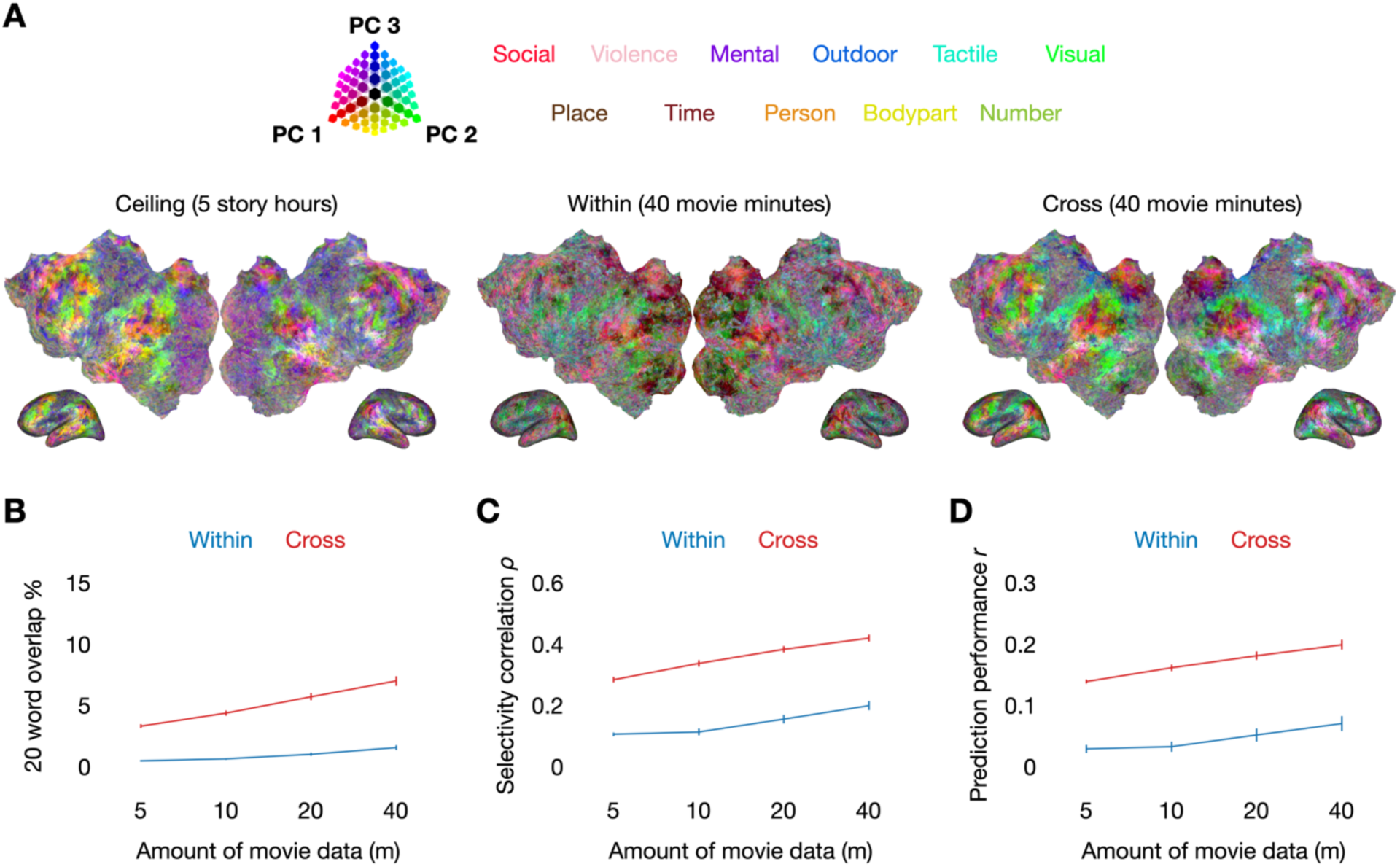
Rapid cross-modal semantic mapping. Cross-participant and within-participant encoding models were trained on brain responses to silent movies. To train within-participant encoding models on brain responses to movies, semantic features were extracted from official audio descriptions. (**A**) Semantic selectivity was visualized by projecting encoding model weights onto the first 3 principal components of a group space. A cross-participant model trained on 40 minutes of movie responses from a goal participant was able to capture the similar patterns of semantic selectivity as a ceiling encoding model trained on 5 hours of story responses from the goal participant. Differences in visual cortex and auditory cortex were caused by the difference in modality. (**B**) Semantic selectivity was operationalized by simulating responses to 10,470 words and identifying the 20 words with the highest simulated responses. Cross-participant models had higher word overlap with the ceiling model than within-participant models trained on the same amount of data from the goal participant. (**C**) Semantic selectivity was alternatively operationalized by simulating responses to 10,470 words and ranking the words based on the simulated responses. Cross-participant models had higher selectivity correlation with the ceiling model than within-participant models trained on the same amount of data from the goal participant. (**D**) Prediction performance was quantified using the linear correlation between predicted and actual response time-courses for a held-out test story. Cross-participant models had higher prediction performance than within-participant models trained on the same amount of data from the goal participant.

Comparing quantitative metrics, we found that cross-participant models had higher word overlap with the ceiling model (**Figure 4B**), higher selectivity correlation with the ceiling model (**Figure 4C**), and higher prediction performance (**Figure 4D**) than within-participant models trained on the same amount of data from the goal participant (*q*(FDR) < 0.05; two-sided paired t-test).

Unlike for story-based transfer, the advantage of cross-participant modeling did not decrease as we increased the amount of movie data from the goal participant. This may reflect the fact that these within-participant models rely on semantic features extracted from verbal descriptions, which do not fully capture the richness of the movies.

### 3.4. Rapid auditory mapping

While the previous analyses mapped semantic representations, naturalistic stories provide access to many other types of linguistic representations (Hamilton & Huth, 2018). Here we assessed whether cross-participant modeling can also be used to map lower-level linguistic representations of sounds and phonemes. We trained reference encoding models by representing the story stimuli using two lower-level feature spaces, and used the same approach as before to transfer the encoding models to the goal participant.

We modeled acoustic information by extracting spectrograms from audio waveforms of the stimulus stories (de Heer et al., 2017; LeBel et al., 2021). To obtain selectivity estimates, we used the encoding models to simulate brain responses to 3,838 sound clips from an external set of narrative stories. To evaluate the selectivity estimates of the cross-participant and within-participant models, we computed the rank correlation with the selectivity estimates of the ceiling model. We visualized the difference in selectivity correlation between cross-participant and within-participant models trained on 24 minutes of data from the goal participant (**Figure 5A**). The cross-participant model had a higher selectivity correlation in auditory cortex, which indicates that it provides more accurate selectivity estimates. Across cortex, cross-participant models had higher overall selectivity correlation (**Figure 5B**) and prediction performance (**Figure 5C**) than within-participant models trained on the same amount of data from the goal participant (*q*(FDR) < 0.05; two-sided paired t-test), but the advantage of cross-participant modeling decreased as we increased the amount of data from the goal participant.

**Figure 5.**
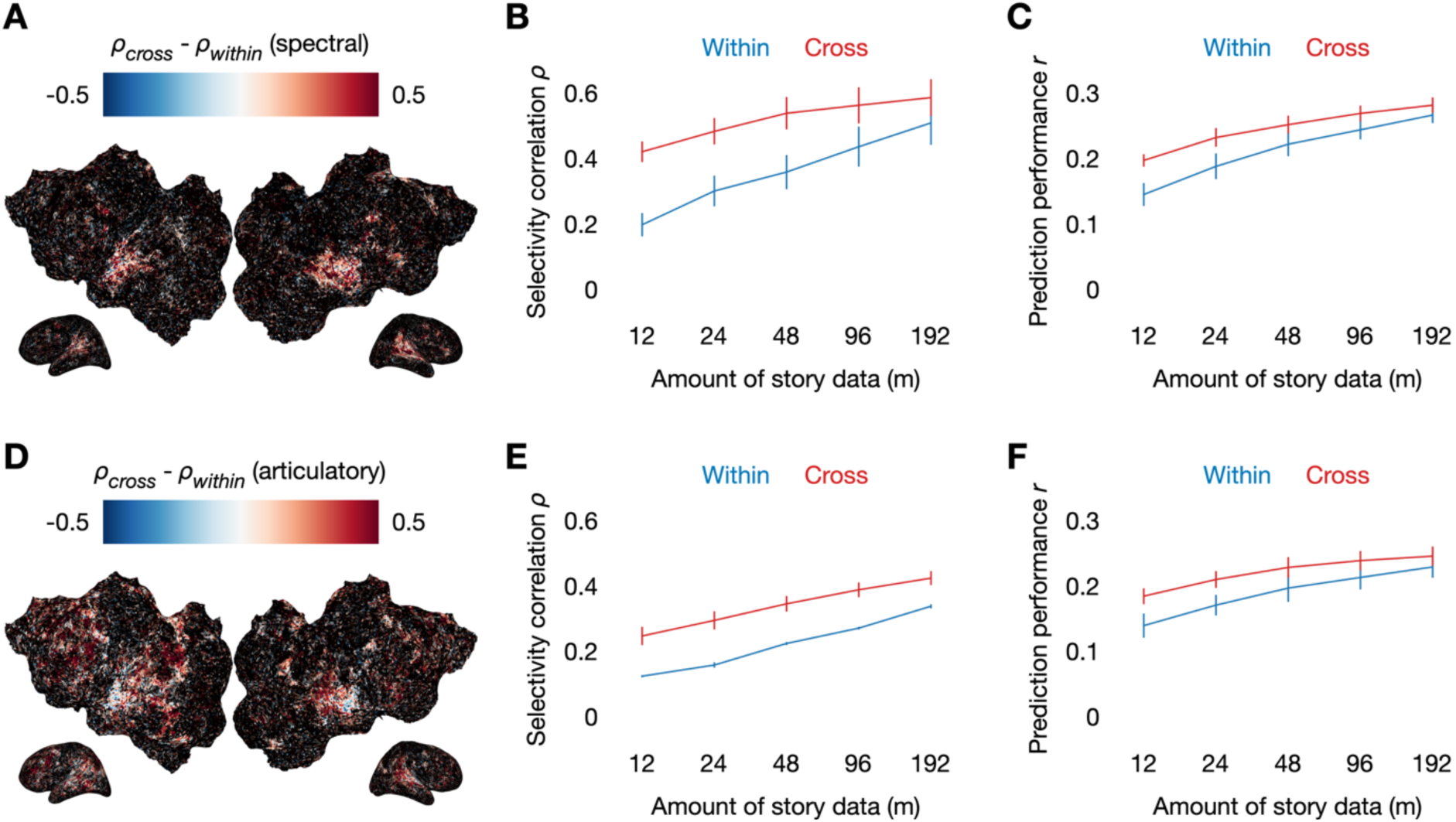
Rapid auditory mapping. (**A**) Acoustic encoding models were trained to predict brain responses based on 256 bands of a mel-frequency spectrogram. Selectivity was operationalized by simulating responses to 3,838 sound clips and ranking the clips based on the simulated responses. Cross-participant models trained on 24 minutes of data from the goal participant had higher selectivity correlation with the ceiling model than within-participant models trained on 24 minutes of data from the goal participant. (**B**) The advantage of cross-participant modeling decreased as the models were trained on more data from the goal participant. (**C**) Cross- participant models had higher prediction performance than within-participant models trained on the same amount of data from the goal participant, but the advantage decreased as the models were trained on more data from the goal participant. (**D**) Articulatory encoding models were trained to predict brain responses based on place of articulation, manner of articulation, voice, and tongue position features. Selectivity was operationalized by simulating responses to 39 phonemes and ranking the phonemes based on the simulated responses. Cross-participant models trained on 24 minutes of data from the goal participant had higher selectivity correlation with the ceiling model than within-participant models trained on 24 minutes of data from the goal participant. (**E**) The advantage of cross-participant modeling decreased as the models were trained on more data from the goal participant. (**F**) Cross-participant models had higher prediction performance than within-participant models trained on the same amount of data from the goal participant, but the advantage decreased as the models were trained on more data from the goal participant.

We modeled articulatory information by extracting the place of articulation, manner of articulation, voicing, and tongue position for each phoneme in the stimulus stories. Previous studies have found that articulatory representations are shared across speech perception and production (D’Ausilio et al., 2009; Watkins et al., 2003; Wilson et al., 2004), so mapping articulatory representations during speech perception could be useful for localizing implants for brain-computer interfaces (Prakash et al., 2025; Verwoert et al., 2025). To obtain selectivity estimates, we used the encoding models to simulate brain responses to 39 phonemes. To evaluate the selectivity estimates of the cross-participant and within-participant models, we computed the rank correlation with the selectivity estimates of the ceiling model. We visualized the difference in selectivity correlation between cross-participant and within-participant models trained on 24 minutes of data from the goal participant (**Figure 5D**). The cross-participant model had a higher selectivity correlation in multiple cortical regions, which indicates that it provides more accurate selectivity estimates. However, the cross-participant model provided less of an advantage in early auditory cortex, which may reflect the fact that the within-participant model is already effective at estimating the selectivity of this region even with limited data. Across cortex, cross- participant models had higher selectivity correlation (**Figure 5E**) and prediction performance (**Figure 5F**) than within-participant models trained on the same amount of data from the goal participant (*q*(FDR) < 0.05; two-sided paired t-test), but the advantage of cross-participant modeling decreased as we increased the amount of data from the goal participant.

These results demonstrate that the benefits of cross-participant modeling are not limited to mapping semantic representations. As new feature spaces are developed to train increasingly accurate models of the brain, it is plausible that these models can also be transferred across participants.

## 4. Discussion

Our results demonstrate the potential for rapid cortical mapping in new goal participants by leveraging data from previously scanned reference participants. By comparing cross-participant models to ceiling models trained on many hours of data from the goal participants, we found that cross-participant models provide accurate selectivity estimates. By comparing cross-participant models to within-participant models trained on the same amount of data from the goal participants, we found that cross-participant modeling is more effective than within-participant modeling in data-limited settings.

Cross-participant modeling could help enable new clinical applications of fMRI. Most current clinical applications are restricted to mapping a small number of broad regions that can be reliably localized with limited data (Matthews et al., 2006; Silva et al., 2018). Conversely, research studies have used fMRI to produce increasingly detailed maps of complex cognitive processes (Huth et al., 2012, 2016). By leveraging large datasets from participants who are able to undergo many hours of scanning, cross-participant modeling may help bridge the gap between research findings and clinical practice. This could be particularly important for monitoring and treating cognitive disorders, since the brain regions that subserve higher level cognition have traditionally been difficult to map with limited data. Cross-participant modeling does not make any assumptions about anatomy, so reference encoding models can be transferred to a new individual even if the organization of cortical representations is disrupted. Moreover, a separate cross-participant model is independently estimated for each goal voxel, so cortical representations can be mapped in spared regions even if other regions are damaged.

Cross-participant modeling could also enable new paradigms in neuroscience research. Currently, “deep” paradigms use a large amount of data from a small number of participants (Kupers et al., 2024). These paradigms can produce detailed cortical maps and highly performant encoding models for each participant, but it is costly to scan enough participants to assess individual and group level differences. Conversely, “wide” paradigms use a small amount of data from a large number of participants (Marek et al., 2022; Ooi et al., 2025). These studies have more power to study individual and group level differences, but are restricted to relatively simple tasks that can be mapped with limited data. Cross-participant modeling combines the advantages of these paradigms—cortical maps can be defined using a large amount of data from a small number of participants, and then transferred to a large number of participants using a small amount of data. This can enable robust individual and group level comparisons of complex cognitive processes.

There are multiple avenues for improving cross-participant encoding models that do not require collecting more data from the goal participant. Our results indicate that cross-participant encoding models scale with the amount of data from each reference participant and the number of reference participants. It may also be possible to improve cross-participant encoding models by developing specialized functional alignment stimuli or collecting multiple repeats of the functional alignment stimuli from the reference participants. Finally, the cross-participant modeling framework does not rely on any specific alignment algorithm or feature space, so cross-participant models should benefit from future improvements in functional alignment and voxelwise modeling.

## Data and Code Availability

All data used in the analysis will be made publicly available at OpenNeuro before publication. All code used in the analysis will be made publicly available at GitHub before publication.

## Author Contributions

Conceptualization J.T. and A.G.H.; Methodology J.T.; Software and resources J.T.; Investigation and data curation J.T.; Formal analysis and visualization J.T.; Writing (original draft) J.T.; Writing (review and editing) J.T. and A.G.H.

## Funding

This work was supported by the National Institute on Deafness and Other Communication Disorders under awards F32DC022178 (J.T.) and R01DC020088 (A.G.H.).

## Declaration of Competing Interests

A.G.H. and J.T. are inventors on a provisional patent application (the applicant is The University of Texas System Board of Regents) that is directly relevant to the cross-participant mapping approach used in this work.

## Acknowledgements

We thank C. Chen for discussions about this project.

## Appendices

**Figure A1.**
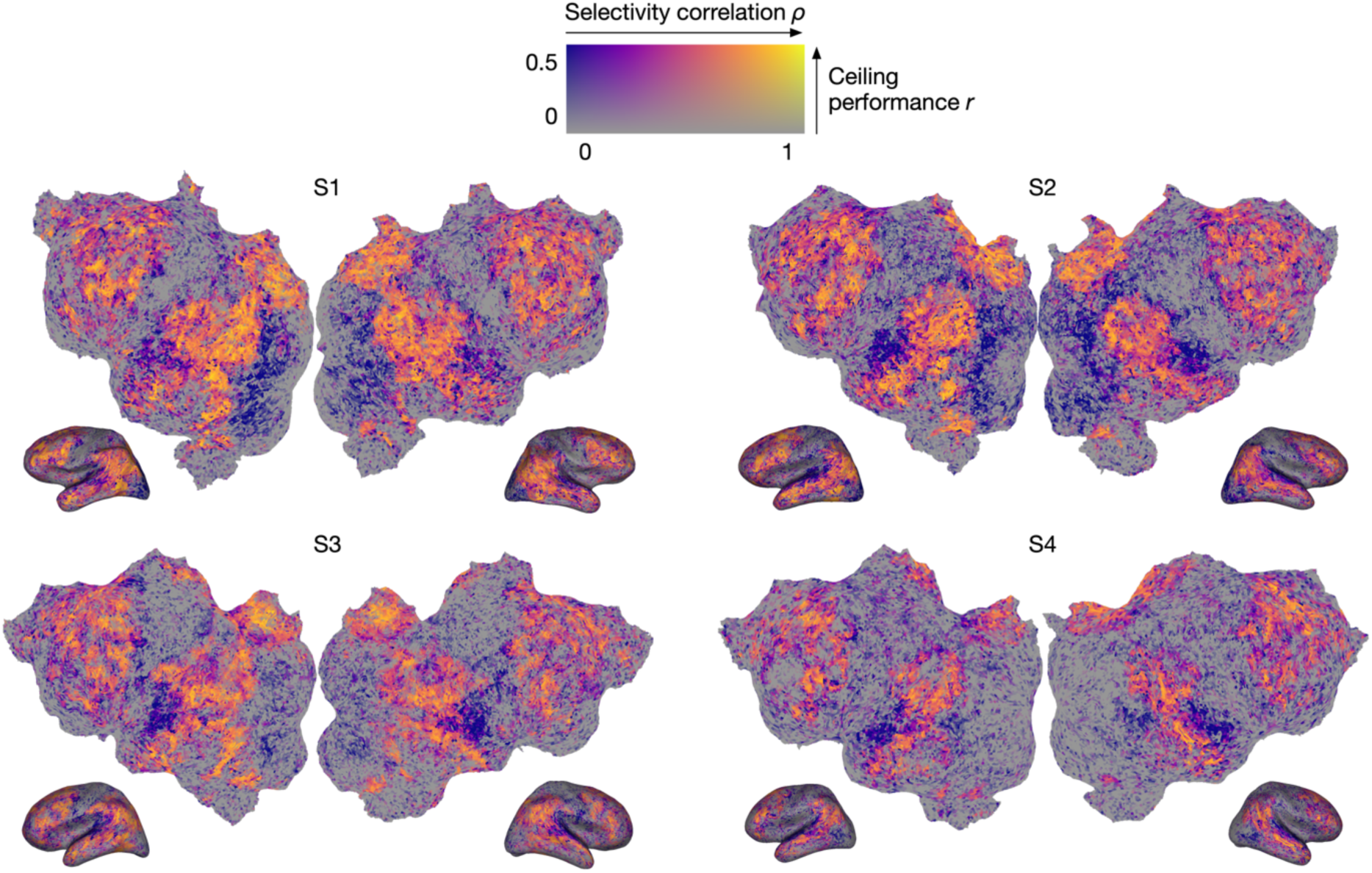
Selectivity correlation for movie-based transfer. Cross-participant encoding models were trained on brain responses to 40 minutes of silent movies. Ceiling encoding models were trained on brain responses to 5 hours of stories. Semantic selectivity was operationalized by simulating responses to 10,470 words and ranking the words based on the simulated responses. Cross-participant models had high selectivity correlation with the ceiling models in most well-predicted regions outside of visual cortex and auditory cortex.

